# FPocketWeb: Protein pocket hunting in a web browser

**DOI:** 10.1101/2022.05.27.493797

**Authors:** Yuri Kochnev, Jacob D. Durrant

## Abstract

**Motivation:** Detecting macromolecular (e.g., protein) cavities where small molecules bind is an early step in computer-aided drug discovery. Multiple pocket-detection algorithms have been developed over the past several decades. Among them, *fpocket*, created by Schmidtke and Le Guilloux, is particularly popular. Like many programs used in computational-biology research, *fpocket* requires users to download and install an executable file. That file must also be run via a command-line interface, further complicating use. An existing *fpocket* server application effectively addresses these challenges, but it requires users to upload their possibly proprietary structures to a third-party server.

**Results:** The FPocketWeb web app builds on this prior work. It runs the *fpocket3* executable entirely in a web browser without requiring installation. The pocket-finding calculations occur on the user’s computer rather than on a remote server.

**Availability:** A working version of the open-source FPocketWeb app can be accessed free of charge from http://durrantlab.com/fpocketweb.

## Introduction

Proteins perform various cellular functions, ranging from mediating cell-signaling pathways to catalyzing vital chemical reactions. Many of these functions depend on interactions between protein receptors and endogenous small-molecule ligands that bind in cavities (pockets) typically located on protein surfaces. Carefully designed exogenous molecules can also bind in protein pockets, competing directly or indirectly with endogenous ligands and thus altering protein activity. Such molecules can serve as useful scientific tools (chemical probes) for revealing the basic biology of proper protein function. When properly optimized, they can also become pharmaceuticals that modulate the activity of disease-implicated proteins.

Structure-based computer-aided drug discovery (CADD) leverages known protein structures to identify and design novel exogenous ligands. Central to these efforts is identifying the protein pocket where the ligand will bind. In many cases, the location of the ligand-binding pocket is well - characterized. For example, crystallographic or NMR structures may include a pocket-bound lig -and, and biochemical experiments can identify pocket-lining catalytic residues that interact with endogenous small-molecule substrates. But often, the location of the ligand-binding pocket is uncharacterized.

In such cases, cavity detection programs can serve as valuable tools for pocket identification. These programs include (a) “geometric” approaches that search for cavities based on the positions of protein-receptor atoms, and (b) “energy-based” approaches that further consider the physicochemical properties of protein residues to identify sites that might energetically favor ligand binding^1^. Among the few programs that fall into the energy-based category, the opensource tool *fpocket* is particularly popular. *fpocket* is written in C, is relatively easy to use, and is notably fast.

As with many CADD programs, the original *fpocket* implementation requires users to (1) download an executable program, (2) choose proper configuration parameters, (3) run the program from a Unix- or DOS-like command-line interface, and (4) separately analyze and visualize the program output. The *fpocket* authors have taken several measures to address these challenges. To facilitate output analysis and visualization, *fpocket* saves output files in formats compatible with VMD^2^ and PyMOL^3^, two popular visualization programs. And to enable use without the command line, the authors created a helpful web server so users can run pocket-hunting calculations in the cloud^4^.

These efforts have substantially improved usability, but some challenges remain. VMD- and PyMOL-formatted output files enable more accessible analysis than text-based output. Still, users must separately download and install these third-party visualization programs, and each has a substantial learning curve. Similarly, the *fpocket* server app eliminates command-line use, but it requires users to upload their (possibly proprietary) structures to a third-party system. Furthermore, in principle, hosting such an app requires an extensive and difficult-to-maintain backend computing infrastructure. Long wait queues may be necessary if a remote resource becomes saturated with requests.

To build on these prior efforts to enhance *fpocket* usability, we created FPocketWeb, a WebAssembly (Wasm) enabled web app that runs *fpocket* entirely in users’ browsers, without the need for other plugins or programs. Wasm-complied code runs locally on the user’s computer, so it do not depend on remote infrastructure to run complex calculations. Instead, a simple web server–or even a “thin server”–is sufficiently powerful to send the Wasm-compiled binaries to the user’s browser.

We release FPocketWeb under the terms of the permissive open-source Apache License, Version 2.0, which is compatible with *fpocket3’s* MIT license. The source code is available for download at https://durrantlab.pitt.edu/fpocketweb-download, without registration. The public browser app is available at http://durrantlab.com/fpocketweb.

## Materials and methods

### Wasm compilation

We downloaded the *fpocket3* codebase, written in the C programming language, from https://github.com/Discngine/fpocket_in_2022. We then used Emscripten 3.1.4 to compile it to a Wasm module. The *fpocket* source code required only minor adjustments to be *Wosm*-compilable. To create the output directories and subdirectories, the original *fpocket* uses a system call to execute the *mkdir* command. This command is readily available on operating systems such as Linux, but it is not available in the browser environment. We therefore commented out the offending line before Wasm compilation. We separately create the required directories on the virtual file system using JavaScript instead.

We also did not implement the full functionality of the VMD molfile plugin^5^. Specifically, our compilation excludes references to the netCDF library, which allows command-line *fpocket3* to load netCDF files. FPocketWeb therefore supports only PDB files.

### FPocketWeb browser app

#### Technical details

The FPocketWeb graphical user interface (GUI) allows users to quickly load protein structures into the browser’s memory, specify FPocketWeb parameters, run calculations using the Wasm-compiled *fpocket* module, and visualize/download the results. To create the interface, we followed the same approach we have used previously,^6–9^. The FPocketWeb interface is written in the Microsoft TypeScript programming language, which compiles to JavaScript. The open-source Vue.js framework (https://vuejs.org/) provides consistently styled components (e.g., buttons, text fields, etc.). Many of these components are derived from Boot-strapVue (https://bootstrap-vue.js.org/), an opensource library that implements the Bootstrap4 framework (https://getbootstrap.com/). We also adapted our existing molecular-visualization Vue.js component^6,8,9^, which leverages the 3Dmol.js library^10^. Finally, we used Webpack (https://webpack.js.org/) and Google Closure Compiler (https://developers.google.com/clo-sure/compiler) to compile, assemble, and optimize the code for size and speed.

#### Usage

##### Input parameters tab

To run FPocketWeb, users need only visit http://durrantlab.com/fpocketweb. The “Input Parameters” tab then appears, which includes the “Input File” and “Advanced Parameters” subsections (Figure 1A and Figure 1B). In the “Input File” subsection (Figure 1A), users can specify the protein file (PDB format) for pocket hunting. This file is opened locally, but it is never uploaded to any third-party server. Users can also click the “Use Example File” button to load *H. sapiens* heat shock protein 90 (Hsp90, PDB 5J2V^11^) for testing. Loaded files are displayed in the “PDB Preview” subsection (not shown) using the 3Dmol.js molecular viewer^10^.

**Figure 1.**
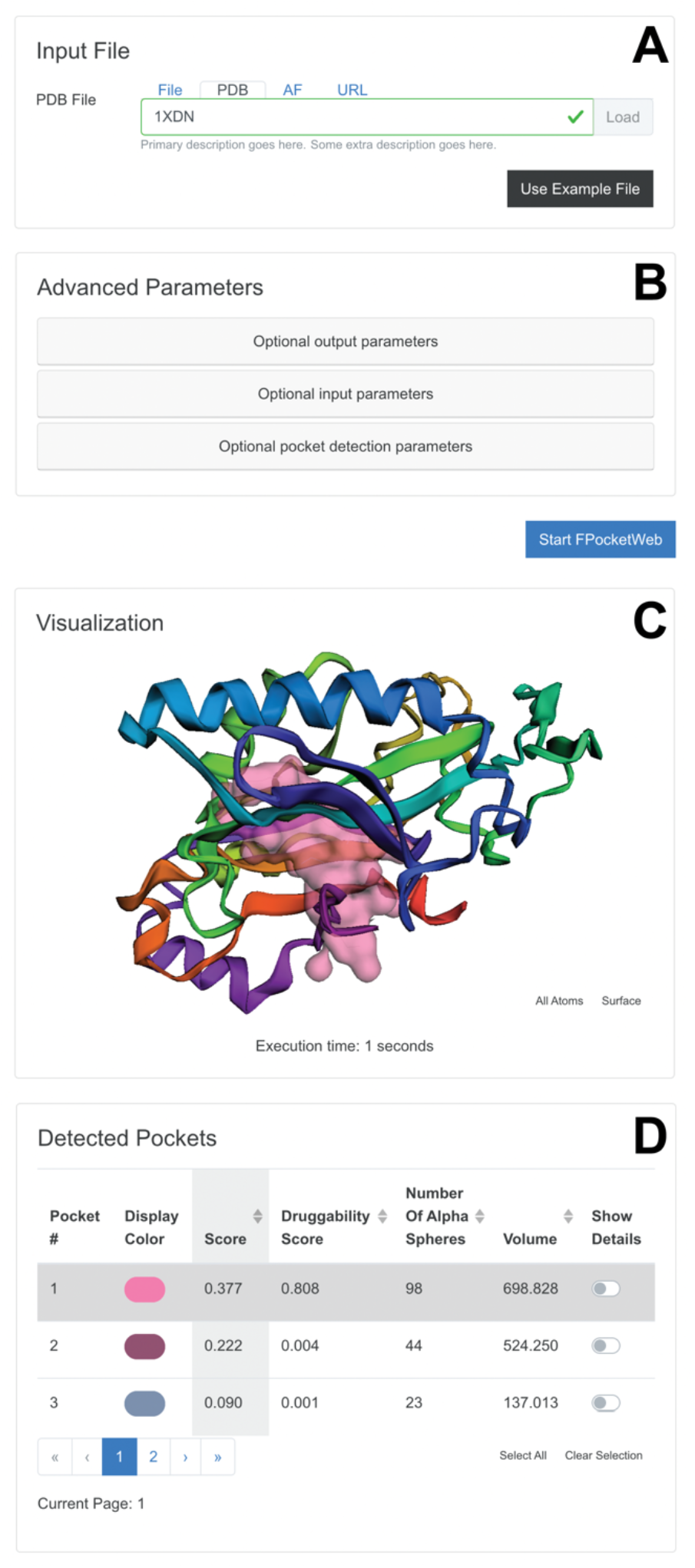
Select subsections of the “Input ? Parameters” and “Output” tabs. (A) The “Input File” _-_ subsection allows users to specify the protein for analysis. (B) The “Advanced Parameters” subsection allows users to fine-tune the pocket-hunting ļ algorithm. (C) The “Visualization” subsection shows ^1^ the detected pockets. (D) The “Deteted Pockets”. subsection describes the properties of each pocket.

The “Advanced Parameters” subsection (Figure 1B) allows users to specify the same parameters available through the *fpocket* command-line executable so they can fine-tune the underlying pocket-finding method. Those interested in further details should consult the original *fpocket* manuscripts^1,4^. FPocketWeb initially hides these parameters because most users will prefer to use the default values. Once ready, users can click the
“Start FPocketWeb” button (Figure 1C) to initiate the FPocketWeb run.

##### Output tab

FPocketWeb displays the “Output” tab once the calculations finish (Figure 1). This tab includes the “Visualization” and “Detected Pockets” subsections. The “Visualization” subsection shows the protein and detected cavities (Figure 1C). Initially, only the pocket with the highest *fpocket* score is shown. The “Detected Pockets” subsection (Figure 1D) contains a table with the detailed output for each detected pocket, one pocket per row. Initially, only the score, the druggability score, the number of alpha spheres, and the volume are displayed, but the “Show Details” toggle allows users to display all other *fpocket* metrics for each detected pocket. Users can also change the color used to visualize each pocket.

The “Output Files” subsection (not shown) allows users to view the FPocketWeb output PDB file, which includes the original protein structure and all the detected pockets. Users can press the “Download” button to save the file. Finally, the “Run Fpocket from the Command Line” subsection (also not shown) provides a code snippet so users can run *fpocket* from the command line with the same FPocketWeb parameters used in the browser.

## Results and discussion

### Benchmarks and compatibility

We performed benchmark calculations on a Linux laptop (HP Pavilion 15.6”) running Fedora 34 (1.3 GHz Intel Core i7 processor and 16 GB 2400 MHz DDR4 memory) to compare FPocketWeb and command-line *fpocket* (Table 1). The two programs had comparable execution times and produced identical scores, druggability scores, numbers of alpha spheres, and volumes. After these initial benchmarks, we further tested FPocketWeb on the browser/operating-system combinations shown in Table 2. The app is compatible with desktop and mobile operating systems, as well as all major browsers (e.g., Chrome, Edge, Safari, and Firefox).

**Table 1.**
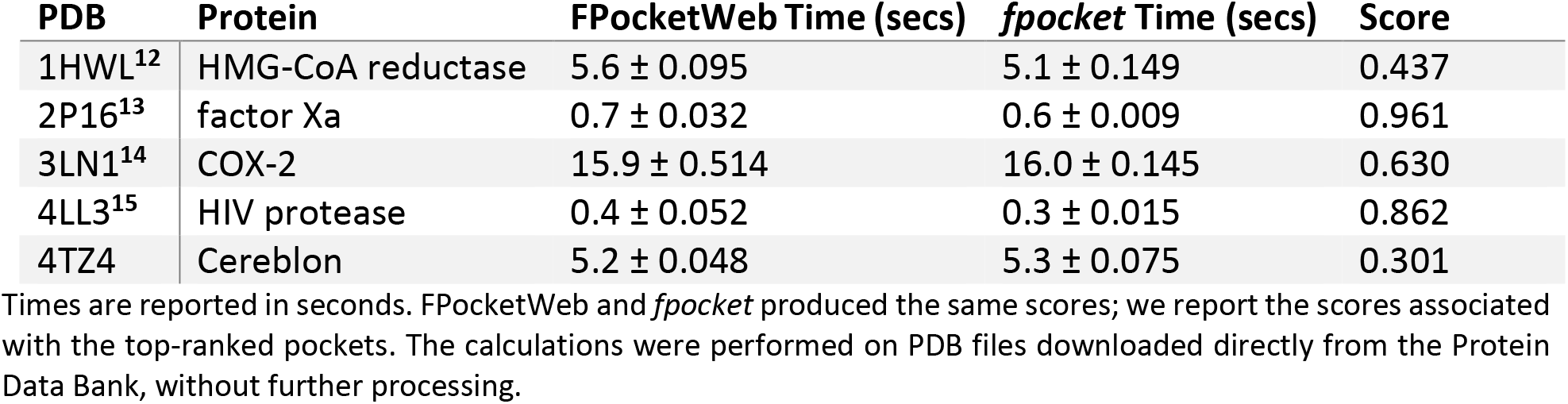
FPocketWeb/*fpocket* benchmarks.

**Table 2.** Browser compatibility.

| Browser | Operating System |
| --- | --- |
| Chrome 101.0.4951.41 | macOS 12.1 |
| Firefox 99.0.1 | macOS 12.1 |
| Safari 15.2 | macOS 12.1 |
| Chrome 101.0.4951.41 | Windows 10 Enterprise |
| Firefox 99.0.1 | Windows 10 Enterprise |
| Edge 100.0.1185.50 | Windows 10 Enterprise |
| Chrome 100.0.4896.127 | Android 12 |
| Firefox 99.2.0 | Android 12 |
| Safari 15 | iOS 15.4.1 |
| Firefox 99.0 | Ubuntu 20.04.4 LTS |
| Chrome 101.0.4951.41 | Ubuntu 20.04.4 LTS |

We have tested FPocketWeb on all major browsers and operating systems.

### Example of use: TEM-1 ß-lactamase

As a demonstration of use, we first considered TEM-1, a bacterial β-lactamase. TEM-1 contributes to penicillin and cephalosporin resistance in Gram-negative bacteria^16^ by hydrolyzing the four-carbon ring common to all β-lactam antibiotics. Aside from the orthosteric pocket where the ring is hydrolyzed, a recently discovered allosteric cryptic pocket^17–20^ is collapsed and hidden (PDB ID: 1FQG^21^) unless co-crystallized with a bound ligand (PDB ID: 1PZP^20^). TEM-1 is often used to benchmark computational tools for cryptic-pocket identification^17,22,23^.

We applied FPocketWeb to TEM-1 crystal structures with (Z)-3-[(4-phenylaza - nylphenyl)amino]-2-(2H-1,2,3,4-tetrazol-5-yl)prop-2-enenitrile (FTA, PDB ID: 1PZP^20^) and penicillin G (open form; PDB ID: 1FQG^21^) bound in the allosteric and orthosteric pockets, respectively.

We first used FPocketWeb to remove all non-protein atoms before detecting pockets on the protein surfaces.

This analysis shows the dramatic impact that the allosteric inhibitor FTA has on the TEM-1 structure. The top-ranked pocket detected when we applied FPocketWeb to the FTA-bound structure (PDB ID: 1PZP^20^) encompassed the allosteric and orthosteric sites (Figure 2A), with a total volume of 2161 Å^3^. In contrast, the allosteric pocket was not detected when we applied FPocketWeb to the penicillin-bound structure (PDB ID: 1FQG^21^) because the allosteric pocket was unoccupied and collapsed. Even the orthosteric pocket ranked only fifth (Figure 2B, 350 Å^3^).

**Figure 2.**
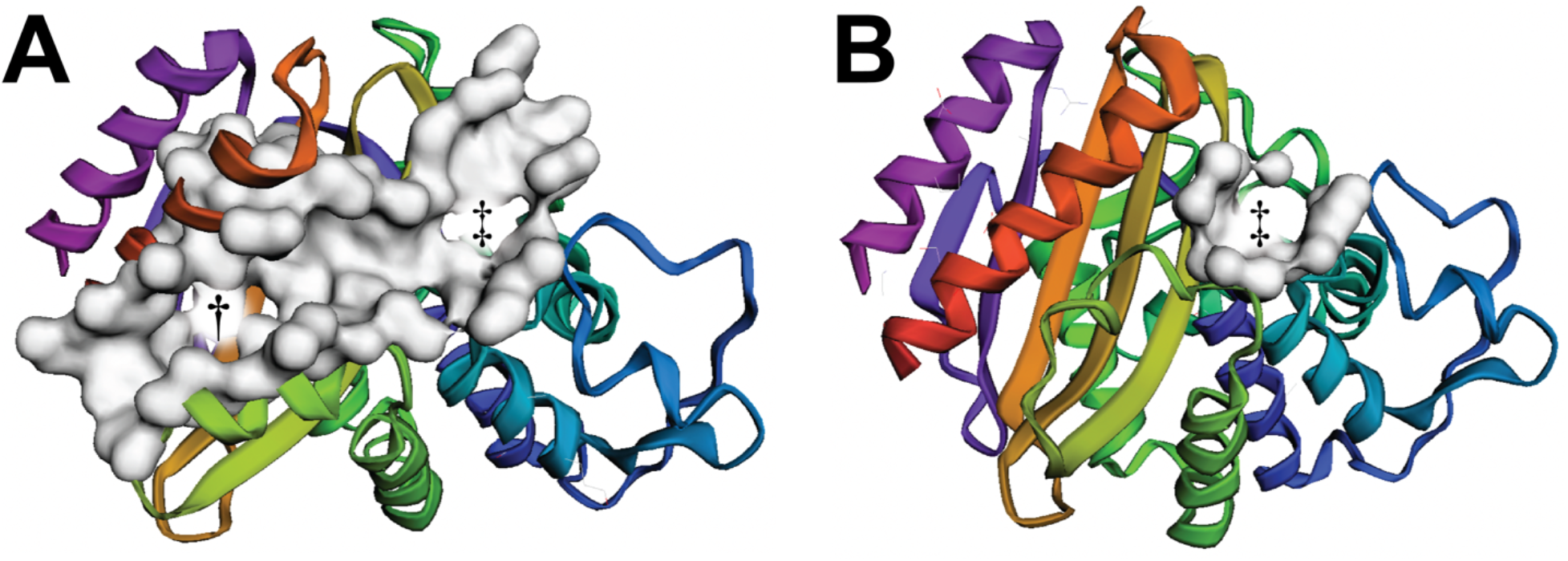
Screenshots of FPocketWeb applied to TEM-1. (A) TEM-1 when the ligand FTA is bound in the allosteric site (PDB ID: 1PZP). The top-ranked pocket, shown in surface representation, encompasses both the allosteric and orthosteric pockets, marked with a dagger and double dagger, respectively. (B) TEM-1 when penicillin G (open form) is bound in the orthosteric site (PDB ID: 1FQG). The fifth-ranked pocket, shown in surface representation, corresponds to the orthosteric pocket (double dagger). The allosteric pocket was not detected because it is collapsed.

### Example of use: influenza neuraminidase

As a second demonstration of use, we consider the enzymatic pocket of influenza neuraminidase (N1). N1 is an exosialidase that prevents the aggregation of newly formed viral particles on infected-cell surfaces^24^. The primary N1 enzymatic pocket^25,26^ is persistent even in the absence of a bound ligand. But a flexible pocket-adjacent “150 loop”^27–29^ enables a pocket extension (the 150-cavity) that is collapsed in many crystal structures. Others have designed N1 inhibitors that exploit this cavity^30^.

We applied FPocketWeb to N1 crystal structures in open (PDB ID: 2HTY^29^) and closed (PDB ID: 2HU4^29^) 150-cavity conformations. We again removed any non-protein atoms before detecting pockets on the protein surfaces. The top-ranked pocket detected when we applied FPocketWeb to the open-150-cavity structure included the 150-cavity (Figure 3A, marked with a dagger and shown as a white surface, 911 Å^3^). The fifth-ranked pocket corresponded to the known sialic-acid binding site (Figure 3Figure 3A, marked with a double dagger and shown as a red surface, 491 Å^3^). In contrast, when we applied FPocketWeb to the closed-150-cavity structure, the 150-cavity was not detected. The top-ranked pocket did include the known sialic-acid binding site (Figure 3B, marked with a double dagger, 1142 Å^3^).

**Figure 3.**
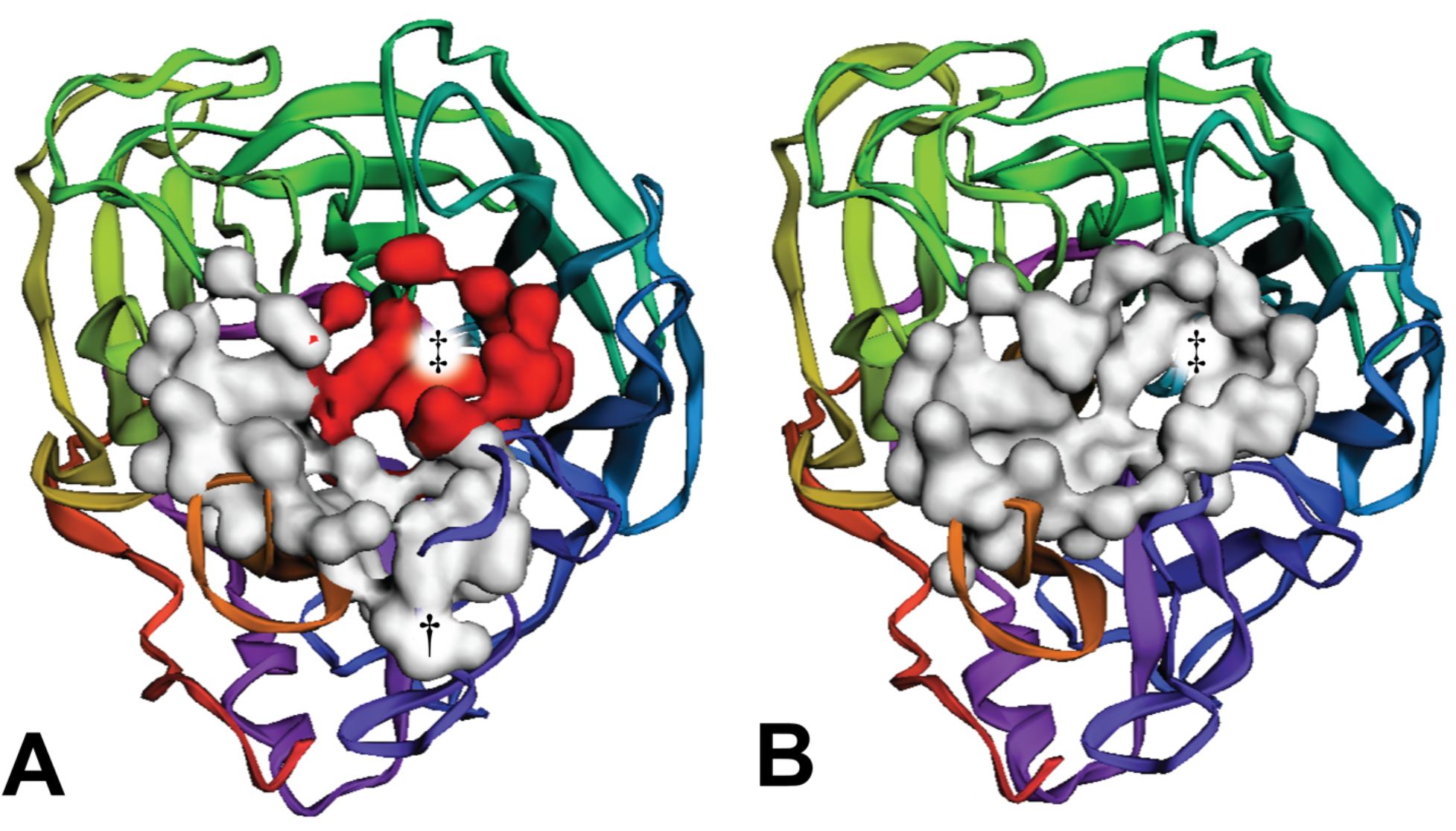
Screenshots of FPocketWeb applied to influenza neuraminidase. (A) Neuraminidase when the 150-cavity is open (PDB ID: 2HTY). The top-ranked pocket, shown in white surface representation, encompasses the 150-cavity (marked with a dagger). The fifth-ranked pocket, shown in red surface representation, corresponds to the sialic-acid binding site (marked with a double dagger). (B) Neuraminidase when the 150-cavity is closed (PDB ID: 2HU4). The top-ranked pocket, shown in white surface representation, encompasses the sialic-acid binding site (marked with a double dagger).

### Limitations

In compiling *fpocket* to Wasm, we also compiled the related programs specified in the same makefile, including *mdpocket* (a tool for analyzing pocket conformations sampled by molecular dynamics simulations), *dpocket* (a tool for extracting physicochemical pocket descriptors), and *tpocket* (a tool for evaluating cavity scoring functions). However, we subsequently focused our web-based implementation and testing on *fpocket* alone. Implementing *mdpocket* as a browser app is also appealing, but it would require loading an entire MD trajectory into the browser’s memory, which is not presently practical. For those interested in using an online version of *mdpocket*, we recommend the original *fpocket* creators’ useful website, which allows users to upload their trajectory files and run *mdpocket* in the cloud^4^.

Another limitation is that FPocketWeb only accepts files in the PDB format. The original *fpocket3* executable uses the molfile plugin^5^ to load structure and trajectory files in many formats. In some cases (e.g., the netCDF format), the molfile plugin leverages other libraries with many dependencies. Given that PDB is the *de facto* standard for static protein structures and that FPocketWeb is not meant to work with trajectory files (i.e., it does not implement *mdpocket),* we opted to focus on the PDB format alone. Users with files in other formats can easily convert them to the PDB format using programs such as VMD^2^ and Open Babel^31^.

### Conclusion

FPocketWeb is an easy-to-use tool that allows users to (1) run *fpocket* entirely in a web browser and (2) analyze/visualize any identified pockets. We release it under the Apache License, Version 2.0. The source code is available free of charge from http://durrantlab.com/fpocketweb-download, and the web app can be accessed at http://durrantlab.com/fpocketweb.

## Acknowledgments

We acknowledge the University of Pittsburgh’s Center for Research Computing for providing valuable computer resources.

